# Genomic characterization of *Francisella tularensis* and other diverse *Francisella* species from complex samples

**DOI:** 10.1101/2022.08.07.503100

**Authors:** David M. Wagner, Dawn N. Birdsell, Ryelan F. McDonough, Roxanne Nottingham, Karisma Kocos, Kimberly Celona, Yasemin Özsürekci, Caroline Öhrman, Linda Karlsson, Kerstin Myrtennäs, Andreas Sjödin, Anders Johansson, Paul S. Keim, Mats Forsman, Jason W. Sahl

## Abstract

*Francisella tularensis*, the bacterium that causes the zoonosis tularemia, and its genetic near neighbor species, can be difficult or impossible to cultivate from complex samples. Thus, there is a lack of genomic information for these species that has, among other things, limited the development of robust detection assays for *F. tularensis* that are both specific and sensitive. The objective of this study was to develop and validate approaches to capture, enrich, sequence, and analyze *Francisella* DNA present in DNA extracts generated from complex samples. RNA capture probes were designed based upon the known pan genome of *F. tularensis* and other diverse species in the family *Francisellaceae*. Probes that targeted genomic regions also present in non-*Francisellaceae* species were excluded, and probes specific to particular *Francisella* species or phylogenetic clades were identified. The capture-enrichment system was then applied to diverse, complex DNA extracts containing low-level *Francisella* DNA, including human clinical tularemia samples, environmental samples (*i.e.*, animal tissue and air filters), and whole ticks/tick cell lines, which was followed by sequencing of the enriched samples. Analysis of the resulting data facilitated rigorous and unambiguous confirmation of the detection of *F. tularensis* or other *Francisella* species in complex samples, identification of mixtures of different *Francisella* species in the same sample, analysis of gene content (*e.g.*, known virulence and antimicrobial resistance loci), and high-resolution whole genome-based genotyping. The benefits of this capture-enrichment system include: even very low target DNA can be amplified; it is culture-independent, reducing exposure for research and/or clinical personnel and allowing genomic information to be obtained from samples that do not yield isolates; and the resulting comprehensive data not only provide robust means to confirm the presence of a target species in a sample, but also can provide data useful for source attribution, which is important from a genomic epidemiology perspective.

## Introduction

The genus *Francisella*, and the family it is assigned to – *Francisellaceae*, include an expanding and diverse group of organisms. The most well-known species is *F. tularensis*, causative agent of the zoonosis tularemia; it was first described in 1912 [1]. Since then, new *Francisella* species have continued to be described (https://lpsn.dsmz.de/genus/francisella), suggesting even more species will be described in the future [2]. The known species exhibit diverse lifestyles, including endosymbionts (*Francisella*-like endosymbionts, FLEs), pathogens of humans and other animals, and free-living environmental species. Despite these diverse lifestyles, there is significant genomic overlap among *Francisella* species [3].

Genomically characterizing new *Francisella* species, including their genomic overlap with *F. tularensis*, is important for efforts to accurately detect and identify *F. tularensis*. *F. tularensis* is a highly monomorphic species [4] and, as such, it is not difficult to identify genomic targets conserved across all representatives of this species (*i.e.*, its core genome) and, thus, avoid false negative results by using detection assays targeting the core genome. However, almost all of the *F. tularensis* core genome is also found in other *Francisella* species [2]. As such, the main challenge with designing DNA-based detection assays for *F. tularensis* that are both highly specific and highly sensitive is avoiding false positive results. Indeed, our recent extensive analysis of all available genome sequences from *F. tularensis* and all other *Francisellaceae* species (*i.e.*, *F. tularensis* genetic near neighbor species) revealed just six unique coding region sequences (CDSs) within the *F. tularensis* core genome that are not found in any isolated near neighbor species [2]. This pattern of reduced genomic space available for species-specific detection/diagnostics due to overlap with near neighbor species, which we previously demonstrated with *Burkholderia pseudomallei* and described as signature erosion [5], becomes more extreme as additional near neighbor species are discovered and incorporated into signature erosion analyses.

A significant challenge to accurate signature erosion analysis for *F. tularensis* is that many *Francisellaceae* species can be difficult or impossible to culture. Indeed, environmental surveys followed by gene-based DNA analyses (*e.g.,* 16S rRNA genes, succinate dehydrogenase [*sdhA*]) have revealed the presence of multiple diverse, previously unknown *Francisella* species [6]. In some cases, use of novel culturing methods has led to the successful isolation of *Francisella* species from environmental samples [7, 8], allowing for the subsequent generation of whole genome sequence (WGS) data and description of these previously unknown species [9, 10]. However, this is not the case for all unknown *Francisella* species and these undescribed near neighbor species are thought to be the source of many false positive results generated by some diagnostic platforms focused on detection of *F. tularensis* DNA from air filter samples [11]. In addition, even if a species (*e.g.*, *F. tularensis*) can be isolated from some sample types, this is not the case for all sample types; for example, human samples containing *F. tularensis* DNA that are collected after the administration of antibiotics or air filters containing *Francisella* DNA but no live organisms.

To address this challenge, we developed a DNA capture and enrichment system based upon the collective pan genome of all known *Francisellaceae* species. We demonstrate that this system, when coupled with subsequent sequencing of the enriched DNA, generates robust genomic data for diverse *Francisella* species from a variety of complex sample types, including human clinical samples, environmental samples, and air filter samples. Analysis of the resulting data facilitates rigorous and unambiguous confirmation of the detection of *F. tularensis* and other *Francisella* species in complex samples, identification of mixtures of different *Francisella* species in the same sample, analysis of gene content, and high-resolution whole genome-based genotyping.

## Materials and Methods

### Bait system design

The pan-genome of a set of 424 *Francisellaceae* genomes (S1 Table) was determined with LS-BSR v1.2.3 [12], resulting in 60,889 unique CDSs. CDSs shorter than 120nts were filtered from the analysis, leaving 50,676 CDSs. CDSs were sliced into 120nt fragments, overlapping by 60nts, resulting in 572,897 potential probes. These probes were clustered with USEARCH v11 [13] at 80% ID, resulting in 368,660 remaining probes. Probes were then screened against *F. tularensis* rRNA genes (5S, 16S, 23S) with LS-BSR, and probes with a blast score ratio (BSR) value >0.8 [14] to any rRNA gene were removed. Remaining probes were aligned back against the set of 424 *Francisellaceae* genomes with LS-BSR and BSR values were calculated. Probes conserved only in a single *Francisellaceae* genome (BSR value >0.8) were removed as they could represent contamination, resulting in a set of 200,053 potential probes. Remaining probes were screened with LS-BSR against a set of >125,000 non-*Francisellaceae* bacterial genomes in GenBank and those with a BSR of >0.8 were removed, resulting in a final set of 188,431 probes (S2 Table). Regions with extremely high GC content (>50% GC) or extremely low GC content (<22%) are considered difficult to hybridize using baits. To increase the likelihood of capturing these regions, our library design included a boosting strategy wherein probes corresponding to these types of regions were multiplied by 2X-10X copies, assigning higher redundancy to the most extreme regions (>70% and <15% GC). The final set of probes was ordered from Agilent. To validate the coverage across a reference *F. tularensis* genome, the live vaccine strain (LVS; GCA_000009245.1), probes were aligned against it with minimap2 v2.22 [15] and the depth of coverage was calculated with Samtools v1.11 [16]. The breadth of coverage was then calculated by identifying the number of bases in the reference genome covered by at least one nucleotide in the probe.

### Samples utilized for DNA capture and enrichment

#### Negative and positive control samples

To evaluate the capture enrichment system, we utilized a complex environmental sample consisting of dust collected in Flagstaff, Arizona as our *Francisella*-negative control (Dust1; S3 Table). DNA was extracted from the dust sample using a Qiagen PowerSoil kit (Qiagen, Hilden, Germany), and 400 ng of the extract (100 uL volume) was spiked with 0.1 ng of genomic DNA obtained from a *F. tularensis* subsp. *tularensis* A.II strain to create a *Francisella*-positive control sample, which was evaluated after both one (Dust2) and two (Dust3) rounds of enrichment.

#### *F. tularensis*-positive clinical tularemia samples from Turkey

These eight clinical samples (F0737, F0738, F0739, F0741, F0742, F0744, F0745, and F0749; S3 Table) were fine-needle lymph node aspirations obtained in 2011 from patients with oropharyngeal tularemia originating from multiple regions of Turkey. As previously described [17], DNA was extracted using a QIAamp DNA Mini Kit (Qiagen, Hilden, Germany) and all resulting DNA extracts were determined to contain *F*. *tularensis* DNA; no *F. tularensis* isolates were obtained from these clinical samples as they were collected after antibiotic therapy had been initiated. The original clinical samples were collected as part of the medical workup for tularemia diagnosis, and residual samples were de-identified and donated for this study. As such, this study does not meet the federal definition of human subjects research according to 45 CFR 46.102 (f) and, therefore, are not subject to Institutional Review Board review.

#### *F. tularensis*-positive animal sample from Arizona

This tissue sample (F1069; S3 Table) was obtained from the spleen of a dead, partially decomposed squirrel [18]; *F. tularensis* was not isolated from this sample. DNA was extracted from the tissue sample using a Qiagen QIAmp kit following the manufacturer’s protocol, which was subsequently confirmed to contain *F. tularensis* DNA [18].

#### *Francisella*-positive air filter samples

These are a subset of a larger set of hydrophobic polytetrafluoroethylene (PTFE, EMD Millipore, Danvers, MA, USA) fluorophore membrane filters (3 μm pore size, 47 mm diameter, 150 μm thickness, and 85% porosity) generated from environmental surveillance efforts conducted in Houston, Texas, USA from 15 June to 22 September 2018, and in Miami, Florida, USA from 30 August to 17 November 2018. Air was sampled above ground using sampling units (PSU-3-H, HI-Q Environmental Products, San Diego, CA, USA) located at multiple permanent locations in Houston and Miami. The specific sampling locations are confidential, but filters collected in the same city on the same day are from different locations. Air was collected onto each filter for 24 hours (± 30 minutes) at an airflow rate of 100 liters per minute (±5%). DNA was extracted from a portion of each collected filter and the resulting DNA extracts were tested for the presence of *Francisella* DNA using one or more PCR assays; details of these initial extraction and PCR methods are confidential. These initial testing procedures identified 43 filter extracts putatively containing *Francisella* DNA (Air1-Air43; S3 Table).

Remaining portions of these 43 filters were provided to NAU and DNA was extracted using Qiagen DNeasy PowerWater Kits following the manufacturer’s protocol with several modifications. Prior to use, precipitation in all solutions was dissolve by either pre-warming to 55°C for 5–10 minutes (reagents PW1 and PW3) or shaking (PW4); PW1 was used while warm to facilitate DNA elution from filters. To facilitate release of DNA from filters, they were subjected to two rounds of vigorous bead beating using customized beads not included in the extraction kit. Each filter portion was rolled with the top side of the membrane facing inward into a sterile 2 mL tube containing customized beads composed of 1.0 g of 0.1 mm zirconia/silica beads together with 0.5 g of 1.0 mm zirconia/beads. These tubes were then incubated at 65°C overnight (12-18 hours) in 1 ml of PW1 solution and then homogenized using a FastPrep 24TM 5G Bead Beater (MP BioMedicals, Irvine, CA) using the following settings: manual program, 6.0 m/s, 60s cycles, Matrix C, Volume 1, unit ml, 7 cycles total, 300s rest between each cycle. The tubes were then centrifuged at ≤4000 x g for 1 min at room temperature and the supernatant transferred into 1.5 mL Lo-Bind tube. The tube containing the filter was then refilled with 650 μL warmed PW1 solution and heated at 65°C for 10 minutes prior to repeating the bead beating protocol. To remove all residual debris, the supernatant of DNA eluted from the two cycles of bead beatings was centrifuge at 13,000 x g for 1 min at room temperature and transferred to a clean 1.5 mL Lo-Bind tube (Thermo Fisher Scientific, Waltham, MA). The DNAs were captured using a silica filter column, cleaned, and eluted following the manufacturer’s protocol. All work was conducted in a clean biosafety cabinet and the tools/equipment were decontaminated prior to each use.

#### Tick samples containing FLEs

These five samples were DNA extractions of whole ticks or tick cell lines containing FLEs (S3 Table). Sample D.v.0160 was extracted from an adult female *Dermacentor variabilis* tick collected in Morden, Manitoba, Canada on 17 April 2010; and sample D.v.0228 was extracted from an adult female *D. variabilis* collected in Windsor, Ontario, Canada on 29 May 2011. As previously described [19], DNA was extracted from these whole ticks using a 96-well DNeasy Tissue Kit (Qiagen, Valencia, CA, USA) modified for use with the QIAvac vacuum filtration system, and presence of FLE DNA in the resulting extracts was determined via PCR and targeted gene sequencing. Samples D14IT15.2 and D14IT20 were extracted from two separate *Hyalomma rufipes* ticks both collected on the island of Capri, Italy on 1 May 2014 [20]. As previously described, DNA was extracted from these surface-sterilized whole ticks and detection of *Francisella* DNA in the extracts was determined by PCR [20]. Sample DALBE3 is a tick cell line of *Dermacentor albipictus* that was previously documented to contain an FLE [21]; it was created by Policastro *et al* [22] from ticks collected in Minnesota in 1989 (U. Munderloh, personal communication). This cell line was purchased in 2016 from The Tick Cell Biobank [23] as 2 ml growing cultures in L-15B300 medium [24]. Upon receipt, total DNA was extracted directly from vials using an EZ1 extraction kit (Qiagen, Hilden, Germany). The resulting data from these samples were compared to whole genome sequences previously generated for FLEs present in individual representatives of the tick species *Argus arboreus*, which has been described as *Francisella persica* [25], and *Amblyomma maculatum* [3; ID: FLE_Am]; two representatives of *Ornithodoros moubata*, one generated by Gerhart *et al* [26; ID: FLE_Om] and the other generated by Duron *et al* [27; ID: FLE-Om], two representatives of *Hyalomma asiaticum* [28; IDs: NMGha432, XJHA498, 29], and three representatives of *Hyalomma marginatum* [30; IDs: FLE-Hmar-ES, FLE-Hmar-IL, FLE-Hmar-IT].

### Library preparation

DNA extracts at starting concentrations ranging from 5ng-200ng total were subsequently sheared using QSonica Q800 Sonicator (QSonica, Newtown, CT) at a protocol of 60% amplitude, 15 sec on/off. The final size of ∼250 bp was assessed using a Fragment Analyzer genomic DNA analysis Kit (Agilent Technologies, Santa Clara, CA). For library preparation, a SureSelect XT-low input sample kit reagent (Agilent Technologies, Santa Clara, CA) was used. Briefly, end repair and A-tailing were performed on sheared DNA and an adaptor was ligated to the A-tailed ends of the DNA fragments, followed by purification. Each ligated fragment was uniquely indexed via PCR-amplification for 9 cycles (2 min at 98°C, 9 cycles for 30 s at 98°C, 30 s at 60°C, 1 min at 72°C, and a final extension of 5 min at 72°C), followed by purification. All DNA purification steps were carried out using Agencourt AMPure XP beads (1X bead ratio; Beckman Coulter Genomics, Brea, CA). Library quantity was assessed by Qubit Br dsDNA (Thermo Fisher Scientific, Waltham, MA). The size and quality were assessed an Agilent DNF-374 fragment analyzer. To achieve a final yield of 2000-2500ng per sample, libraries were re-amplified in four replicates for 10-12 cycles and all replicates were pooled.

### Probe-hybridization

To prevent dissociation of probes with AT-rich sequences, we utilized a slow-hybridization protocol involving hybridization reagents from a SureSelect XT0 kit (Agilent Technologies, Santa Clara, CA). Following the manufacturer’s protocol, ∼2000 ng from each library was hybridized with the probes at 65°C for 16-24 hours. Hybridized libraries were recovered using 50 μL of Dynabeads from the MyOne Streptavidin T1 Kit (Thermo Fisher Scientific, Waltham, MA), which were then washed three times using SureSelect wash buffers. We then PCR-amplified directly from the beads using the SureSelectXT-LI Primer Mix (Agilent Technologies, Santa Clara, CA), using the same PCR conditions described above for library preparation except with a 14-cycle parameter to increase the concentration of the capture library. The amplicons were then separated from the beads via a magnetic plate and transferred to a new tube. To amplify the residual capture library remaining on the beads, the beads were used directly for PCR using a KAPA HiFi PCR ready mix (Roche KAPA Biosystems, Wilmington). The captured libraries from both PCR events were then combined and purified. To further enrich for *Francisella* DNA present in the original samples, we performed a second round of enrichment on the captured libraries following the same procedures used for the first enrichment. Final enriched library quantity was assessed by Qubit Br dsDNA (Thermo Fisher Scientific, Waltham, MA), and size and quality were assessed by Fragment Analyzer.

### Sequencing

Sequence libraries were created from enriched DNA using a 500 bp insert and standard PCR library amplification (KAPA Biosystems). Paired end sequences were obtained on either the Illumina Miseq or Nextseq platforms.

### Bioinformatic analyses

#### Read classifications

Reads were mapped against the standard Kraken database with Kraken v2.1.2 [31] and the number of reads classified as *Francisellaceae* was determined. The percentage of *Francisellaceae* reads was determined by dividing the number of Kraken2 *Francisellaceae* classifications by the total number of reads per sample.

#### Read mapping and breadth of coverage calculation

Reads were aligned against the reference genome LVS with minimap2 and the depth of coverage was called with Samtools after filtering read duplicates. The breadth of coverage at a minimum depth of 3x was then calculated with a Samtools wrapper script (https://gist.github.com/jasonsahl/b5d56c16b04f7cc3bd3c32e22922125f). The presence of probes was calculated using the same workflow, but using the probe set as the reference genome.

#### Enrichment metagenome assembly and binning

Enriched reads were assembled with meta-SPAdes v3.13.0 [32] using default settings (--meta) and contigs shorter than 1000 nucleotides were removed. Genomes were binned with concoct v1.1.0 [33] using BAM files created by mapping Illumina reads back against the metagenome assembly with minimap2. Ribosomal RNA genes were extracted from metagenomes with barrnap v0.9 (https://github.com/tseemann/barrnap). Metagenome assembled genomes (MAGs) associated with *Francisellaceae* were combined with reference genomes and a cluster dendrogram was created with MASH distances [34] and Scikit-Bio (http://scikit-bio.org) using a custom script (https://gist.github.com/jasonsahl/24c7cb0fb78b4769521752193a43b219).

#### SNP discovery and construction of phylogenies

SNPs were identified among publicly available genomes, as well as enriched samples assigned to the same species/taxonomic group, by aligning reads against reference genomes using minimap2 v2.22 and calling SNPs from the BAM file with GATK v4.2.2 [35]. Maximum likelihood phylogenies were inferred on the concatenated SNP alignments for each species/taxonomic group using RAxML-NG v0.9.0 [36]. All of these methods were wrapped by NASP [37].

#### *Francisella* Pathogenicity Island (FPI) and antimicrobial resistance (AMR) screening

The genes associated with the FPI were screened in all complex enriched samples containing *F. tularensis* and FLEs by calculating a breadth of coverage at 3x depth. The SNPs associated with rifampicin and streptomycin resistance in *F. tularensis* [38] were screened in all complex enriched samples containing *F. tularensis* with NASP and manually investigated from the resulting SNP matrix.

#### *sdhA* gene analysis

*In silico* PCR (isPCR) was performed with USEARCH (-search_PCR- strand both maxdiffs 2) on metagenomes using primers (5’-AAGATATATCAACGAGCKTTT-3’, 5’-AAAGCAAGACCCATACCATC-3’) previously published for the *sdhA* gene of *Francisella* [6]. The predicted amplicons from metagenomes were added to those extracted from 152 reference *Francisella* genomes (S1 Table) using the isPCR approach. Sequences were oriented to the same strand based on blastn v2.11.0+ [39] alignments and aligned with MUSCLE v5.0 [40]. A phylogenetic tree was inferred on the alignment with IQ-TREE v2.0.3 [41] and the integrated ModelFinder method [42]. isPCR was also performed for two assays, Ft-sp.FTT0376 and Ft-sp.FTS_0772, that were previously identified as highly specific to *F. tularensis* [2].

#### Identification of unique probes

To identify probes in the *F. tularensis* core genome, probes were aligned with BLAT [43] in conjunction with LS-BSR against a set of 327 *F. tularensis* genomes (S1 Table). A probe was considered core if it had a BSR value >0.8 across all target genomes. *F. tularensis* core probes were then aligned against 127 *Francisellaceae* near neighbor genomes (S1 Table) with LS-BSR/BLAT and probes with a BSR value >0.4 in all non-target genomes were discarded. Clade-specific probes were identified for other species and phylogenetic clades using this same approach. Probes conserved in all *Francisellaceae* genomes were identified by a BSR value >0.8 across 498 *Francisellaceae* genomes (S1 Table).

#### FLE annotation

Reads were aligned against *F. persica* ATCC VR-331 (NZ_CP013022) and mutations were called with Snippy v4.6.0 (https://github.com/tseemann/snippy). Mutations were recorded if they resulted in a premature stop or if they resulted in a frameshift mutation.

#### Signature erosion

Genomes were randomly sampled 100 times at different depths from a set of 172 non-*F. tularensis Francisellaceae* genomes (S1 Table) with a custom script (https://gist.github.com/jasonsahl/990d2c56c23bb5c2909d). All *F. tularensis* core genome probes were aligned against the 172 *Francisellaceae* near neighbor genomes with LS-BSR/BLAT and the number of probes with a BSR value <0.4 were identified and plotted.

#### Limit of detection analysis with *WG-FAST*

NASP was run on a set of 327 *F. tularensis* genomes (S1 Table), resulting in a set of 15,805 SNPs. A maximum likelihood phylogeny was inferred on the concatenated SNP alignment with RAxML-NG v0.9.0 [36], which also provided the required files for the whole genome focused array SNP typing (*WG-FAST*) v1.2 tool [44]. Reads from the eight clinical tularemia samples from Turkey were randomly sampled at various depths (1000-10000) using seqtk v1.3 (https://github.com/lh3/seqtk) and processed with *WG-FAST* at a minimum depth of 3x.

## Results and Discussion

### Composition of the bait enrichment system

Probes were designed against the pan-genome of a diverse set of *Francisellaceae* species and specific probes were identified for most species/clades included in the design (S4 Table); these specific probes represent markers that can distinguish between mixtures of different species/clades in complex samples. For *F. tularensis*, 68 specific probes were identified based on conservative homology thresholds. Our previous study identified six CDSs that were specific to and conserved across *F. tularensis* [2]. Of the 68 *F. tularensis*-specific probes in this study, 19 were associated with these previously described CDSs and at least one of these 19 probes mapped to each of the six CDSs; the other 49 probes mapped to other regions in the *F. tularensis* core genome. The probes utilized in this study are only 120 nucleotides in length, whereas CDSs can be much longer, which likely explains the higher number of *F. tularensis*-specific probes compared to specific CDSs. An alignment of all probes in the system (n=188,431) against the *F. tularensis* reference LVS genome demonstrated that ∼72% of the genome was covered using default mapping parameters in minimap2. Because our design clustered probes at 80% identity, the relatively low coverage across LVS compared to the enrichment results described below is likely due to selection of some representative probes from non-*F. tularensis Francisellaceae* species that fail to align across species using default and conservative mapping parameters.

### Signature erosion

We observed significant signature erosion with the *F. tularensis* probes by including near neighbor genomes (Fig 1). When only 100 non-*F. tularensis Francisellaceae* genomes were randomly sub-sampled 100 times from a set of 172 total genomes and analyzed together with 327 *F. tularensis* genomes, a range of 69-253 *F. tularensis*-specific probes were identified, more than the 68 *F. tularensis*-specific probes discovered when using the full set of 172 non-*F. tularensis Francisellaceae* genomes. This result demonstrates the importance of thoroughly sampling genomic near neighbors before designing and testing diagnostics and may partially explain false positive results with some assays used to detect *F. tularensis* [11]. Although we identified 68 probes as specific to *F. tularensis* using the *Francisellaceae* genomes analyzed in this study, not all these 68 probes may be actually specific to *F. tularensis*, as the genomic regions they target could be conserved in other, currently unknown *Francisellaceae* species in the environment. Thus, the number of *F. tularensis*-specific probes may be reduced from 68 as new *Francisellaceae* genomic information becomes available but should still allow for multiple redundant probes specific to *F. tularensis*.

**Fig 1.**
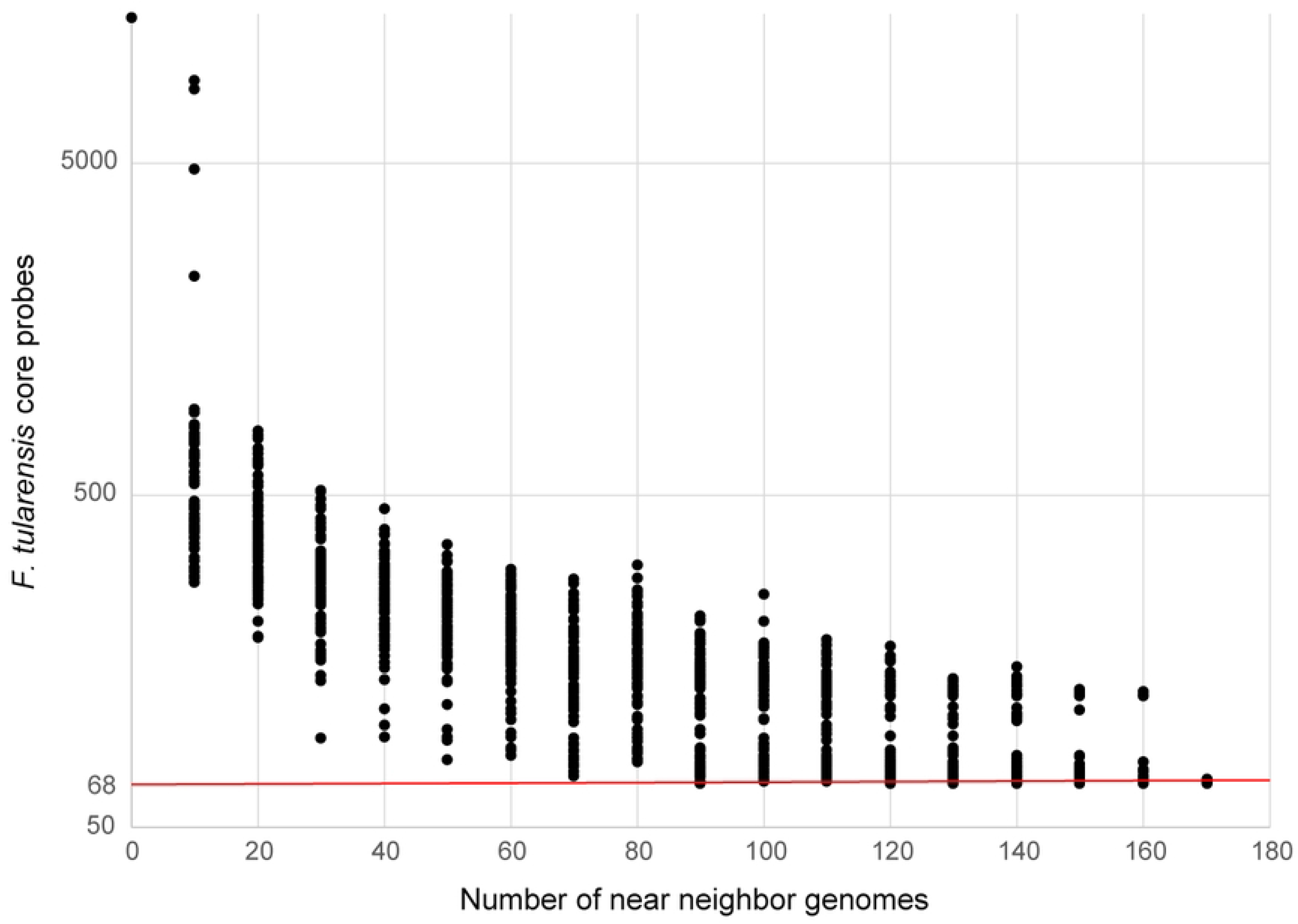
Signature erosion analysis of *F. tularensis*-specific probes. *F. tularensis* near neighbor genomes were randomly sampled 100 times at different depths (0-170, sampled every 10) and then screened with *F. tularensis* core probes (n=5,606) using LS-BSR/BLAT. The number of probes with a BSR value <0.4 were plotted, indicating that they were specific to *F. tularensis* and not present in sampled near neighbor genomes. The red line indicates the number of *F. tularensis*-specific probes (n=68) identified using the entire near neighbor dataset (n=172).

#### Negative and positive control samples

Targeted DNA capture and enrichment greatly increased the proportion of *F. tularensis* DNA present in the dust sample spiked with *F. tularensis* DNA (S1 Fig), which, upon sequencing, generated high breadth of coverage across the *F. tularensis* genome, including across the 68 *F. tularensis*-specific probes (S4 Table), as well as the genomic regions targeted by the two *F. tularensis*-specific PCR assays (S3 Table). Prior to enrichment, the estimated proportion of *Francisellaceae* signal in the spiked dust sample was ∼0.03%. However, after two rounds of enrichment, the signal increased to 69% based on whole genome sequencing data (S1 Fig, S3 Table), resulting in a >10^3^-fold increase in the proportion of *F. tularensis* DNA in the final sample while simultaneously decreasing the non-target background DNA. When reads from the twice enriched Dust3 sample were mapped to the LVS reference genome, 89% of the chromosome was covered at a minimum depth of 3x (S3 Table). This is greater than the 72% coverage just using probes, as described above, likely because no reads were rejected due to mapping across species and/or this region was unintentionally enriched as “by-catch” [45] because it is located adjacent to other *Francisellaceae* genomic regions that were included in the probe design. All *F. tularensis*-specific probes were covered at >80% breadth of coverage in the Dust3 sample based on a 3x depth of coverage; no reads mapped to the 68 *F. tularensis*-specific probes from the sequence data generated from the original spiked dust sample (Dust1; S4 Table). These findings demonstrate that *F. tularensis* genomic DNA present at very low levels in complex samples can be captured, enriched, sequenced, and analyzed using our bait-capture system.

#### *F. tularensis*-positive clinical tularemia samples from Turkey

Sequence reads from all eight of these samples were assembled and binned into one *Francisella* MAG per sample, and also mapped to the genomic regions targeted by the two *F. tularensis*-specific PCR assays and all 68 *F. tularensis*-specific probes, which were covered at >80% by ≥3 reads (S4 Table). There was extensive coverage of all FPI genes in these eight enriched samples (S5 Table), and the SNPs previously associated with rifampicin and streptomycin resistance in *F. tularensis* [38] were not present in these eight samples. These findings demonstrate that the content of specific genes and SNPs of interest can be determined in enriched complex samples containing *F. tularensis*.

Of the total sequencing reads from these enriched samples, 84.2-98.1% mapped to *Francisellaceae*, resulting in ≥3x coverage across >99% of the *F. tularensis* LVS reference genome (S4 Table). Full length 16S rRNA gene sequences (1523 nt) were obtained from five of these samples, despite the removal of probes aligning to the *rrn* operon in the bait design phase. This was likely due to this region being present in these samples in sufficient quantity that it was detectable in these samples via sequencing even without enrichment, and/or by-catch. Within both the global *Francisellaceae* dendrogram (S2 Fig) and the *F. tularensis*-specific ML phylogeny used for analysis of these samples (Fig 2), all eight samples from Turkey were assigned to the major clade corresponding to *F. tularensis* subsp. *holarctica* (Type B), the only subspecies known to occur in Turkey [17, 46] and, within the Type B clade, all of these clinical samples were most closely related to previous isolates obtained from similar geographic regions of Turkey.

**Fig 2.**
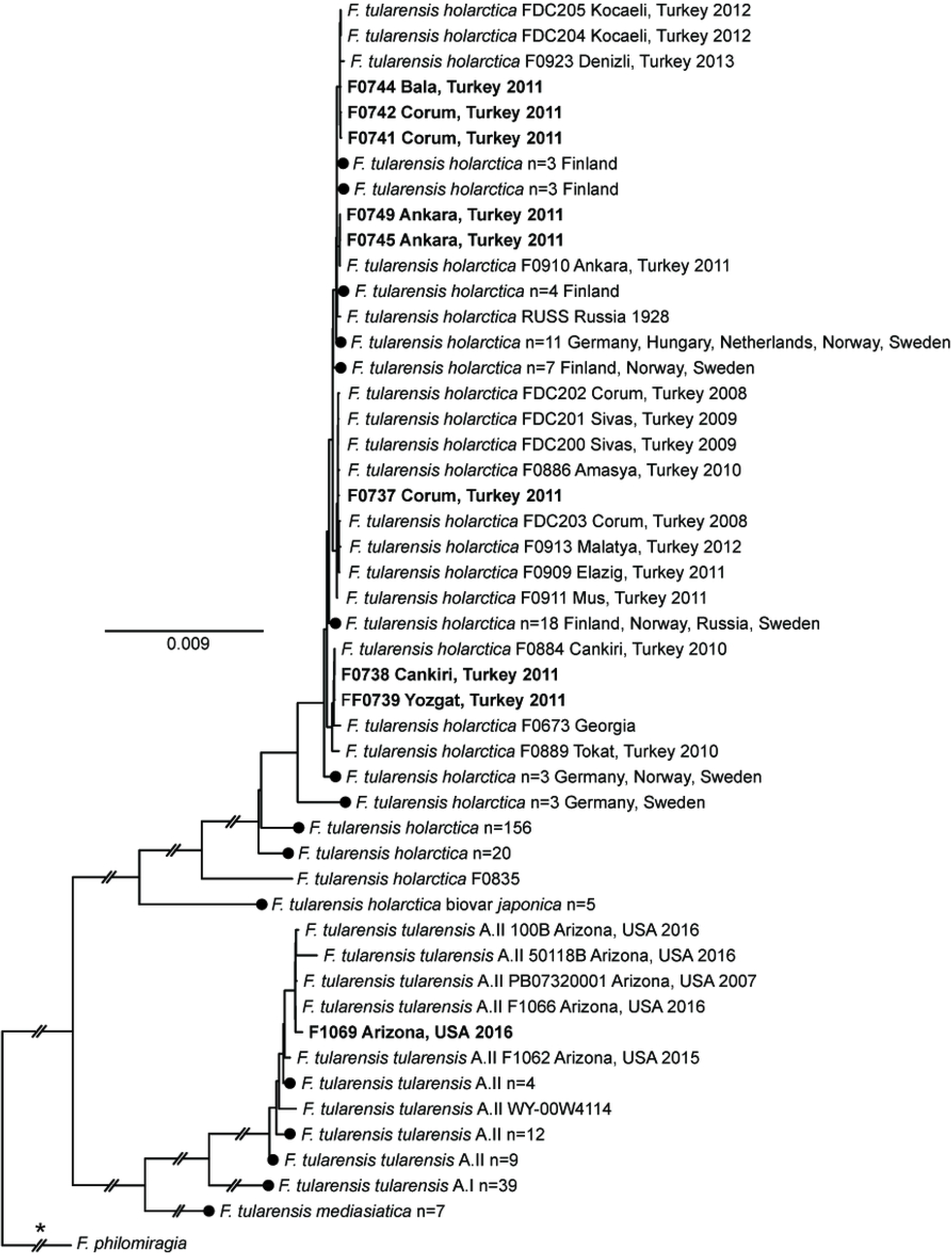
*F. tularensis* phylogeny. Maximum-likelihood phylogeny based upon 15,526 core genome SNPs shared in 327 publicly available *F. tularensis* genomes (S1 Table) and DNA enriched from nine complex samples (bold text). The phylogeny is rooted on the branch indicated with * and some branch lengths have been shortened (double hash marks). Scale bar units are average nucleotide substitutions per site. Black circles indicated collapsed nodes.

Enriched clinical samples F0738 and F0739 were originally obtained in 2011 from individuals residing in Cankiri and Yozgat, Turkey, respectively (S3 Table); these two cities are located ∼130 km apart in north-central Turkey. Within the *F. tularensis* ML phylogeny (Fig 2), F0738 is separated from *F. tularensis* isolate F0884 (alternative ID: F015) by two SNPs; F0884 was obtained from a human throat swab in Cankiri in 2010 [46]. F0739 is closely related to F0738 and F0884, differing from them by just two SNPs; one of these SNPs is autapomorphic in F0739. F0738, F0739, and F0884 group together in a larger clade with F0673, which was isolated from an unknown source in eastern Georgia at an unknown date [47], as well as F0889 (alternative ID: F039), which was isolated from a human in Tokat, Turkey in 2010 [46]; Tokat also is located in north-central Turkey ∼155 km northeast of Yozgat.

Enriched clinical samples F0741, F0742, and F0744 were all obtained in 2011 (S3 Table): F0741 and F0742 from individuals residing in Corum, and F0744 from an individual residing in Bala, which is in Ankara province in central Turkey. These three enriched samples group together in the *F. tularensis* ML phylogeny (Fig 2) with isolates FDC204, FDC205, and F0923. FDC204 and FDC205 were obtained from Kocaeli in northwestern Turkey in 2012 from the throat swab of a human and a contaminated water source that was likely the source of the human infection, respectively [8]. F0923 was obtained in 2013 from water in Denizli in southwestern Turkey [46]. There are two, one, and two autapomorphic SNPs in F0741, F0742, F0744, respectively, that are unique to each of these enriched samples, as well as two synapomorphic SNPs shared by only F0741 and F0742 (Fig 2).

Enriched clinical sample F0737 was originally obtained in 2011 from an individual residing in Corum, Turkey (S3 Table). This sample groups together most closely in the *F. tularensis* ML phylogeny (Fig 2) with isolate F0886 (alternative ID: F027) but separately from enriched clinical samples F0741 and F0742, which also originated from individuals residing in Corum (S3 Table); finding distinct genotypes in the same geographic area is consistent with oropharyngeal human tularemia in Turkey being caused by multiple distinct phylogenetic groups of *F. tularensis* subsp. *holarctica* [8, 17, 46]. F0866 was obtained in 2010 from Amasya in north-central Turkey from the throat swab of a human [46]; Amasya is located ∼80 km northeast of Corum. There are two autapomorphic SNPs in F0737 that are unique to this sample, as well as one synapomorphic SNP shared by F0737 and F0886 (Fig 2). In the *F. tularensis* ML phylogeny (Fig 2), F0737 and F0866 group together in a larger clade with seven other isolates from Turkey.

Enriched clinical samples F0745 and F0749 were obtained in 2011 from two different individuals residing in Ankara, Turkey (S3 Table). These two enriched samples group together in the *F. tularensis* ML phylogeny (Fig 2) and are closely related to *F. tularensis* isolate F0910 (alternative ID: F244), which also was obtained from Ankara in 2011 from the spleen of a rodent [46]. There is one autapomorphic SNP in F0745 that is unique to this sample, as well as one synapomorphic SNP shared by F0745 and F0749 (Fig 2).

Previous attempts to generate sequence data for these eight clinical tularemia samples via metagenomics sequencing of the same DNA extracts used in this study yielded a total of 787,568,687 sequencing reads. However, only 8,848 of those reads (0.001%) assigned to *F. tularensis*, which did not facilitate robust genomic comparisons [48]. In contrast, two rounds of targeted DNA capture and enrichment followed by sequencing –even a minimum of 1000 paired-end reads (see below) – provided robust data to conduct detailed phylogenetic analyses of these same samples, identify novel SNPs within and among them, and examine loci associated with virulence and AMR.

Notably, the previously published *F. tularensis* isolates most closely related to the eight enriched samples also were from Turkey and from the same specific regions from which the patients that yielded the eight enriched samples resided. This suggest the possibility of assigning *F. tularensis* isolates or samples of unknown origin to likely geographic sources via whole genome-based phylogenetic analyses. Of course, the phylogenetic and geographic resolution of such analyses is dependent upon the size and representation of the genomic database to which genomic data from unknown samples/isolates are compared [49]. We utilized the genomic database compiled by Ohrman *et al* [2], which only contains *F. tularensis* isolates that have been whole genome sequenced and includes numerous representatives from Turkey. The enrichment approaches described here will allow the expansion of genomic databases for *F. tularensis* because these databases can now also be populated with genomic data from samples that contain *F. tularensis* DNA but do not yield *F. tularensis* isolates, which will lead to larger databases and facilitate more accurate assignment of unknown isolates to likely geographic sources in the future.

#### *F. tularensis*-positive animal sample from Arizona

Despite being collected from a partially decomposed carcass, the enriched complex sample extracted from the spleen of a dead squirrel (F1069) yielded high-quality *F. tularensis* sequence data. After two rounds of enrichment, 92.73% of the total sequencing reads were classified as *Francisellaceae* (S3 Table), resulting in ≥3x coverage across >99% of the *F. tularensis* LVS reference genome, including the 68 *F. tularensis*-specific probes (S4 Table) and the genomic regions targeted by the two *F. tularensis*-specific PCR assays (S3 Table). This broad and comprehensive coverage of the *F. tularensis* genome allowed the sequencing reads to be assembled into a MAG (1.8Mb) and facilitated detailed phylogenetic analysis of the resulting sample. There was complete coverage of all FPI genes in this enriched sample (S5 Table), and the SNPs previously associated with rifampicin and streptomycin resistance in *F. tularensis* [38] were not present.

Enriched sample F1069 was assigned to the *F. tularensis* subsp. *tularensis* group A.II clade in both the global *Francisellaceae* dendrogram (S2 Fig) and the *F. tularensis*-specific ML phylogeny (Fig 2) and, within that clade, grouped together with F1066; F1069 and F1066 group together in a larger clade in the ML phylogeny (Fig 2) with three other isolates from Arizona. F1069 and F1066 share one synapomorphic SNP and there is one autapomorphic SNP unique to enriched sample F1069. Isolate F1066 (alternate ID: AZ00045112) was obtained from a fatal human pneumonic tularemia case that occurred in Coconino County in northern Arizona; this individual was first evaluated on 6 June 2016 and died 11 June 2016 [18, 50]. The dead squirrel that yielded enriched sample F1069 was collected outside the residence of this victim during an environmental survey conducted on 23 June 2016; multiple rabbit carcasses were also observed during this survey, but none were tested for *F. tularensis* due to decomposition. The patient had regular close contact with her pet dog, which was ill in late May 2016 several days after being found with a rabbit carcass in its mouth; the dog subsequently was determined to be seropositive for exposure to *F. tularensis,* but no isolate was obtained from it. Based upon these findings, the dog was suspected as the source of *F. tularensis* inhaled by the victim. Although genotyping assays assigned both the *F. tularensis* DNA present in the squirrel spleen sample (F1069) and the human isolate to *F. tularensis* subsp. *tularensis* group A.II [18], higher resolution genetic analysis of sample F1069 was not previously possible due to the highly degraded nature of the original sample. The whole genome comparisons presented here, facilitated by capture and enrichment of *F. tularensis* DNA present in complex sample F1069, provides additional strong evidence for the previous suggestion [18] that the victim’s dog obtained *F. tularensis* from wildlife near her residence and then transmitted *F. tularensis* to her.

*F. tularensis* grows slowly on most common laboratory media [51] and represents a risk to unvaccinated laboratory staff growing it [52]; for these and other reasons, diagnosis of tularemia is commonly based upon molecular testing of complex samples (*e.g.*, PCR), serological testing, and/or clinical manifestations rather than culture [53, 54]. And even if an *F. tularensis* isolate is obtained in a clinical laboratory, in the US, once definitively identified as *F. tularensis* that isolate is then subject to Select Agent regulations, which stipulate that the isolate must be destroyed or transferred to a facility authorized to possess Select Agents within seven calendar days [55]; most clinical labs are not registered to possess Select Agents and shipping them is very expensive and complex. We were able to generate whole genome sequencing data for *F. tularensis* human isolate F1066 because, once identified as *F. tularensis*, the isolate was transferred from the clinical laboratory to our Select Agent registered facility at Northern Arizona University where we grew the pathogen and extracted genomic DNA. However, because of these regulations, even if obtained, many clinical *F. tularensis* isolates, at least in the US, are destroyed before they can be whole genome sequenced, which has resulted in a dearth of publicly available whole genome sequences for *F*. *tularensis* strains infecting humans in the US. The enrichment approaches described here offer an opportunity to generate genomic data for *F. tularensis* using DNA extracted from complex clinical samples without culturing, which would not be subject to many Select Agent regulations.

#### *Francisella*-positive air filter samples

After two rounds of capture and enrichment, sequencing of the 43 air filter extracts from the US yielded an average of 9,119,398 (range: 4,882,288-15,966,314) sequence reads per sample, with an average of 66.5% of the reads classified as *Francisellaceae* (range: 11.2-95.5%; S3 Table). The two *F. tularensis*-specific PCR assays yielded negative results for these samples when analyzed *in silico* and although all 43 samples yielded *Francisellaceae* reads (S3 Table), there was no or only minimal mapping of reads (*i.e.*, <40x breadth at a depth of 3x for two probes) to the *F. tularensis*-specific probes (S4 Table), confirming that *F. tularensis* was not present in any of the original samples. Although not unexpected as previous analyses putatively identified *Francisella* DNA in other extracts from these same filters (see above), reads from all 43 samples mapped to probes specific to one or more other *Francisellaceae* spp. and/or clades (S4 Table); this was most pronounced in the 37 samples collected in Houston (Air1-37; S3 Table). Among these 37 samples, reads mapping to *F. opportunistica*-specific probes were present in all 37 samples, reads mapping to *F. novicida*-specific probes were present in 30 samples, and reads mapping to *F. philomiragia*-specific probes were present in 18 samples. Among these same samples, *sdhA* gene sequences consistent with *F. opportunistica* and *F. novicida* were present in 16 and 13 samples, respectively; a *sdhA* sequence in sample Air1 was unique to a clade containing known isolates of *F. orientalis/noatunensis/philomiragia*; and a *sdhA* sequence from Air36 was 100% identical to the *sdhA* sequence present in the only known isolate of *F.* sp. LA11-2445 (Fig 3, S3 Table), which was obtained from a cutaneous infection in southern Louisiana in 2011 [56]. Reads from 17 of the 37 enriched samples collected in Houston were grouped into one (16 samples) or two (one sample, Air1) *Francisella* MAGs each, with an average size of 1,820,822 bp (range: 1,501,353-2,348,11) and containing an average of 780 contigs (range: 211-1,531) per MAG (S3 Table). None of the 18 *Francisella* MAGs grouped in the dendrogram (S2 Fig) with *F. tularensis*, which further confirmed that the 17 filter extracts that yielded these MAGs did not contain *F. tularensis*; however, 14 did group with *F. opportunistica* in the dendrogram, three with *F. philomiragia*, and one with *F. novicida*. SNPs discovered from reads generated from these same 17 samples were used to place the *F. opportunistica*, *F. philomiragia*, and *F. novicida* reads sequenced from these samples into separate ML phylogenies, which also include other published genomes from these species (Figs 4-6).

**Fig 3.**
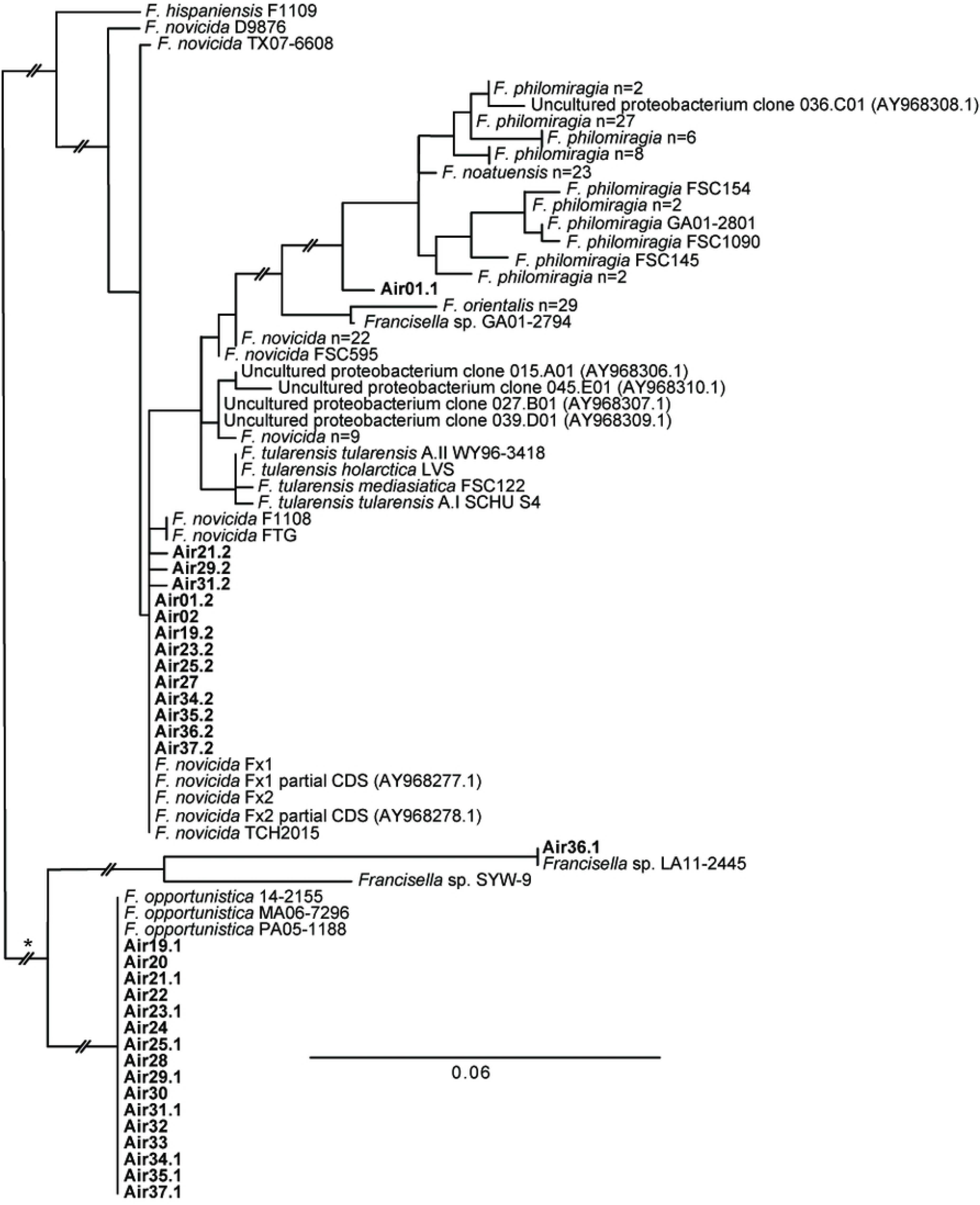
*sdhA* phylogeny. Maximum-likelihood phylogeny inferred from a 348 bp region of the *sdhA* gene extracted from 152 publicly available *Francisella* spp. genomes (S1 Table) and DNA enriched from 20 air filters (bold text), as well as seven publicly available partial gene sequences; accession numbers for the latter are provided in parentheses. Two distinct *sdhA* sequences from the same air filter are indicated as AirX.1/AirX.2. Some strains from the same species with identical *sdhA* genotypes have been collapsed and the number of strains in the collapsed node indicated. The phylogeny is rooted on the branch indicated with * and some branch lengths have been shortened (double hash marks). Scale bar units are average nucleotide substitutions per site.

**Fig 4.**
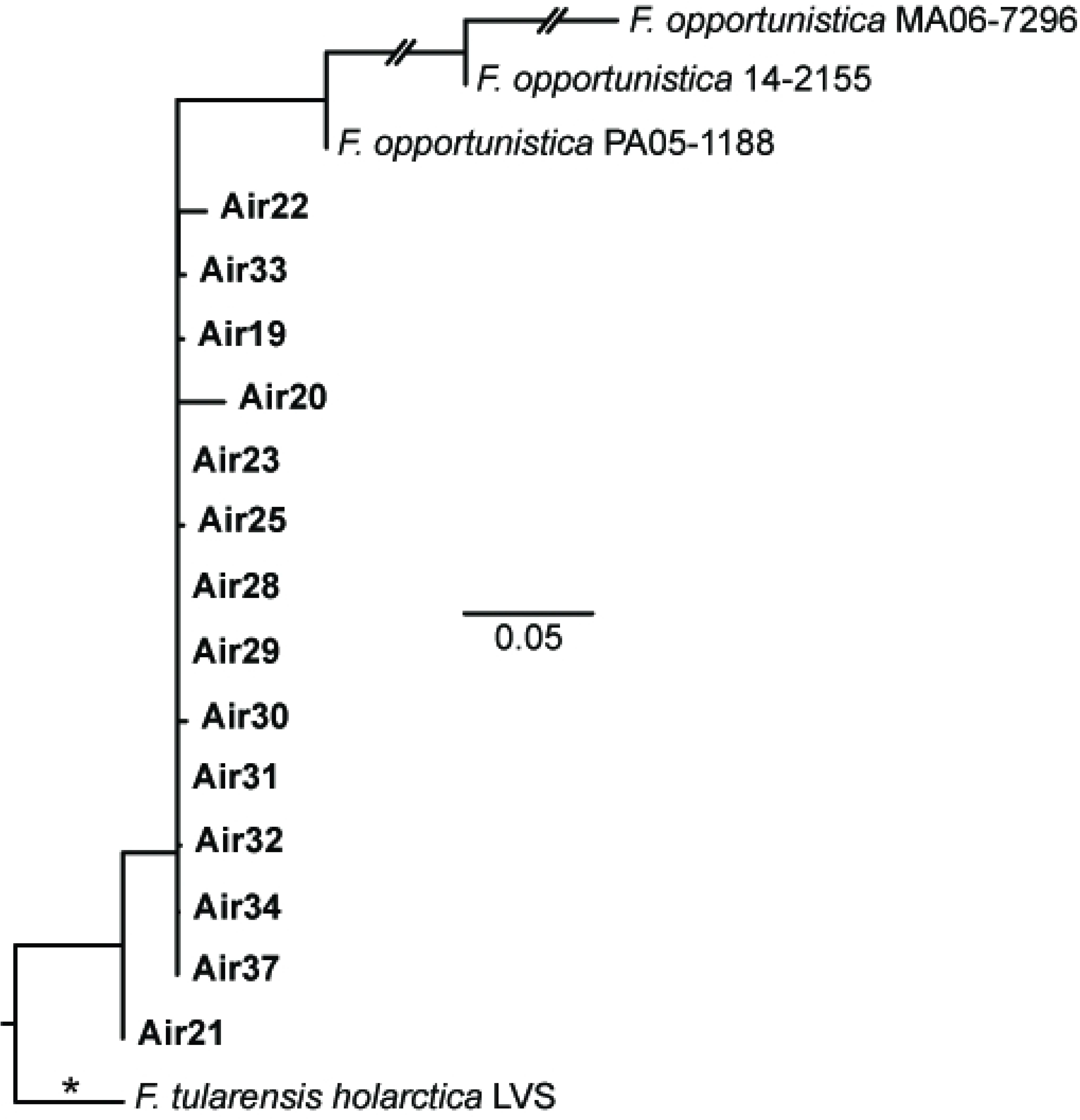
*F. opportunistica* phylogeny. Maximum-likelihood phylogeny based upon 712 core genome SNPs shared in three publicly available *F. opportunistica* genomes (S1 Table) and DNA enriched from 14 air filters (bold text). The phylogeny is rooted on the branch indicated with * and some branch lengths have been shortened (double hash marks). Scale bar units are average nucleotide substitutions per site.

*F. opportunistica sdhA* gene sequences that were 100% identical to each other and to the *sdhA* sequences from the only three known genomes of this species were collected on 16 different air filters in Houston from 21-22 September 2018; a subset of nine of these same air filters also yielded *F. novicida sdhA* sequences (Fig 3). Sequence reads generated from enriched DNA from each of these 16 filters covered >90% of the 2,745 *F. opportunistica*-specific probes (S4 Table), and *Francisella* MAGs identified from sequence data from 14 of these samples (S3 Table) grouped with *F. opportunistica* in the dendrogram (S2 Fig). Interestingly, despite these 14 filters all being collected over just a two-day period, the *F. opportunistica* present on several individual filters differed, as evidenced by unique SNPs for some of these 14 samples in the *F. opportunistica* phylogeny (Fig 4). This pattern, and the collection of these filters at multiple geographic locations, suggest the presence of *F. opportunistica* on these 14 filters was not due to the aerosolization of a single common point source but, rather, the simultaneous aerosolization of multiple unknown sources in the environment, which has also been documented for aerosolized *F. tularensis* [57]. Of note, a storm passed through the Houston area during this time bringing strong winds and 5.9 cm of rain from 20-23 September 2018 (https://www.weather.gov/wrh/Climate?wfo=hgx).

*F. opportunistica* was only recently described as a species [9, 58] and prior to this study was known only from three isolates obtained from immunocompromised humans from different US states in different years. Isolate PA05-1188 was obtained in Pennsylvania in 2005 from an individual with juvenile rheumatoid arthritis and hemophagocytic syndrome, isolate MA06-7296 was obtained in Massachusetts in 2006 from an individual with end stage renal disease [59], and isolate 14-2155 was obtained in Arizona in 2014 from a diabetic individual with renal and cardiopulmonary failure [58]. Based on the locations of these three cases it was previously suggested that *F. opportunistica* may be widespread in the environment [58] but the mechanism by which these individuals became infected with *F. opportunistica* was not clear [59]. Our findings from air-filters in Texas confirm *F. opportunistica* is indeed widespread in the US and suggest that aerosol inhalation should be considered as a route of exposure.

*F. novicida sdhA* gene sequences were present on 13 air filters collected in Houston (Fig 3); these same samples all also yielded sequence reads that mapped to some of the seven *F. novicida*-specific probes (S4 Table). Interestingly, with just two exceptions, the air filters that yielded *F. novicida sdhA* sequences also yielded *sdhA* sequences from other *Francisella* spp. Within the *sdhA* phylogeny (Fig 3), the *F. novicida sdhA* sequences from these 13 air filters were identical or highly similar to each other and to those from three *F. novicida* isolates obtained in or near Houston from humans (Fx1, Fx2, TCH2015) [7, 60, 61], but were more distantly related to four *F. novicida sdhA* sequences (015.A01, 027.B01, 039.D01, 045.E01) generated from soil samples collected in Houston in 2003 [6]. A *F. novicida* MAG was only assembled from sequence reads from one air filter, Air1; a *F. philomiragia* MAG also was assembled from this sample (S3 Table). Within both the MAG dendrogram (S2 Fig) and the *F. novicida* ML phylogeny (Fig 5) the *F. novicida* collected on Air1 is closely related to isolate Fx2.

**Fig 5.**
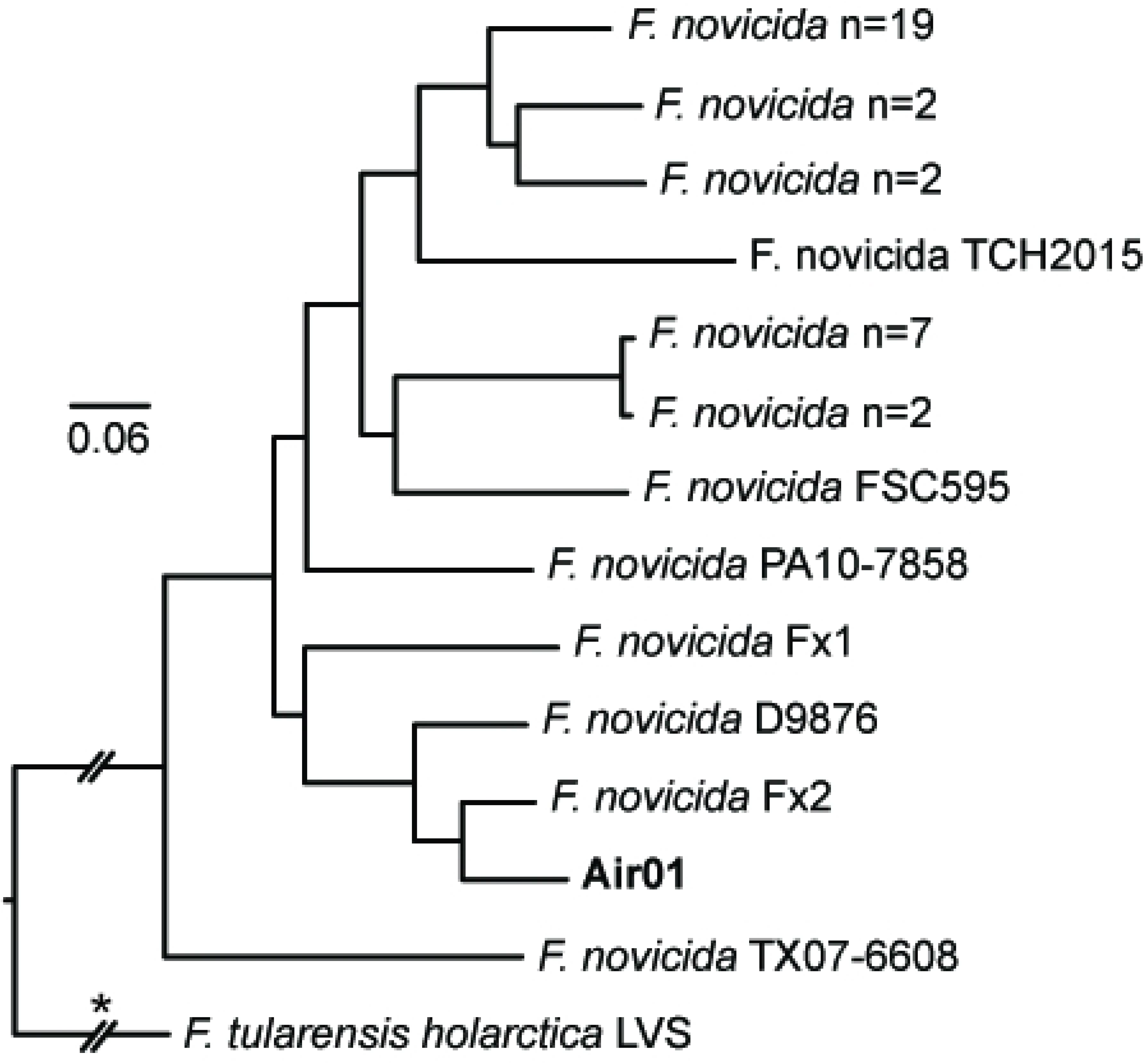
*F. novicida* phylogeny. Maximum-likelihood phylogeny based upon 28,557 core genome SNPs shared in 39 publicly available *F. novicida* genomes (S1 Table) and DNA enriched from one air filter (bold text). The phylogeny is rooted on the branch indicated with * and some branch lengths have been shortened (double hash marks). Scale bar units are average nucleotide substitutions per site.

It is often difficult to determine routes of exposure leading to human infections caused by *F. novicida* and mechanisms of transmission to humans are poorly understood [62]. However, several human infections with *F. novicida*, including the one that yielded isolate Fx2, have presented with pneumonia [60, 63]. This observation, together with our finding that *F. novicida* was aerosolized at multiple locations on multiple dates in Houston in 2018, suggest some human *F. novicida* infections may result from aerosolization and inhalation of this bacterium, like *F. tularensis* [57]. That said, it is important to note that detection of *F. novicida* DNA or DNA from any other *Francisella* spp. on air filters is not evidence that viable bacteria were aerosolized.

*F. philomiragia* MAGs were assembled from sequence reads generated from three air filters – Air1, Air2, and Air12 – collected in Houston on 15, 16, and 21 June 2018, respectively (S3 Table). Sequence reads generated from all three filters mapped to >70% of the 92 *F. philomiragia*-specific probes, and reads from Air1 and Air2 also mapped to >50% of the seven *F. novicida*-specific probes and yielded *F. novicida sdhA* sequences (Fig 3); as described above, Air1 also yielded a separate *F. novicida* MAG. The *F. philomiragia* representatives collected on these filters are basal to all known isolates of this species in both the MAG dendrogram (S2 Fig) and the *F. philomiragia* ML phylogeny (Fig 6) and are quite distinct from each other. Similarly, the *sdhA* gene sequence obtained from Air1 was highly distinct, falling basal to both *F. noatunensis* and *F. philomiragia* (Fig 3), which are sister clades in the global phylogeny of *Francisella* [2], further documenting the distinctiveness of the *F. philomiragia* captured on these filters. The previous *F. philomiragia* isolates included in these analyses were mostly obtained from water or sediment; several were collected in Europe and Asia, but most – and the only known representatives from the US – were obtained in the state of Utah (S1 Table), which is located >1,500 km from Houston.

**Fig 6.**
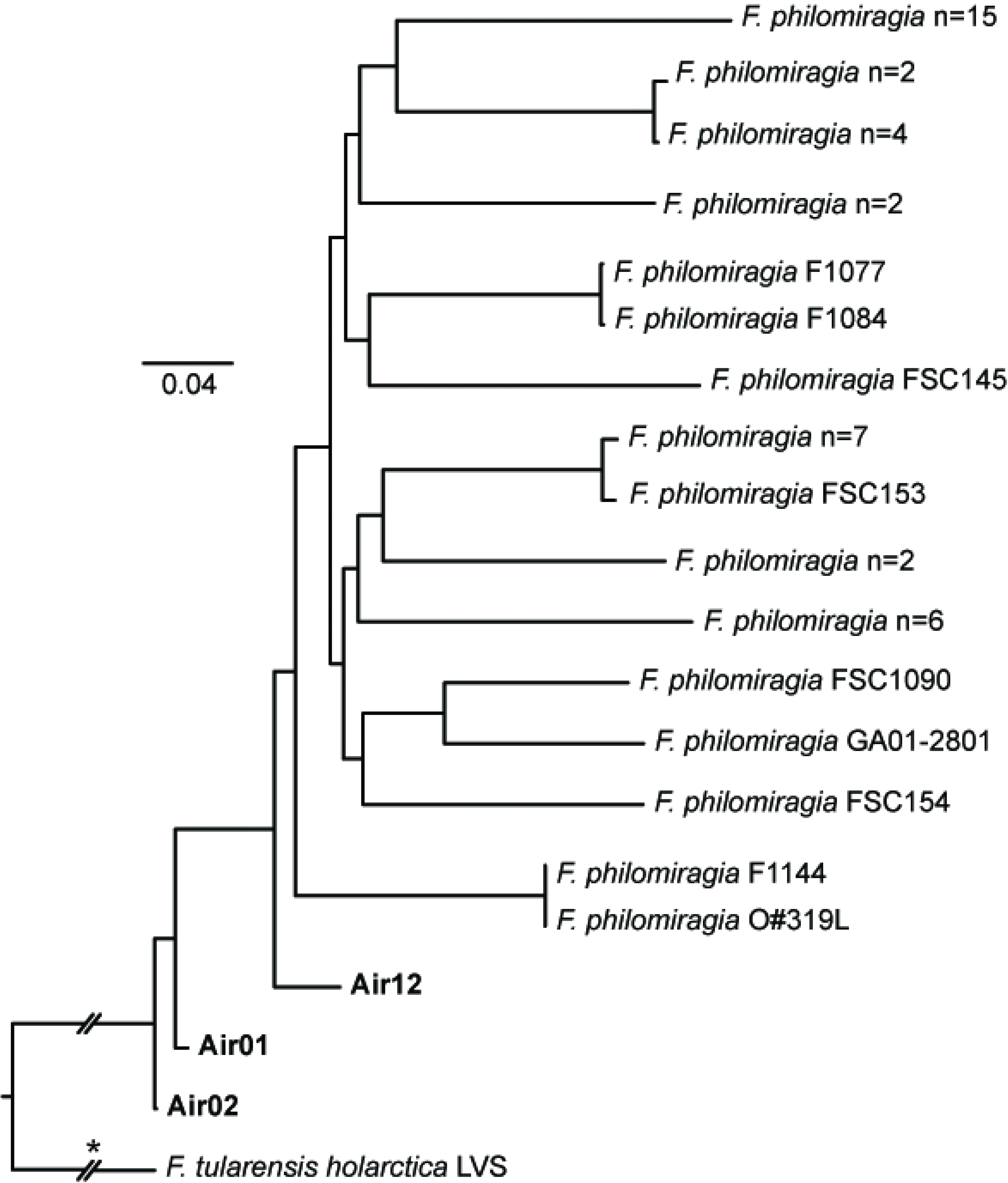
*F. philomiragia* phylogeny. Maximum-likelihood phylogeny based upon 59,753 core genome SNPs shared in 47 publicly available *F. philomiragia* genomes (S1 Table) and DNA enriched from three air filters (bold text). The phylogeny is rooted on the branch indicated with * and some branch lengths have been shortened (double hash marks). Scale bar units are average nucleotide substitutions per site.

*F. philomiragia* is recognized as a rare but serious human pathogen primarily associated with immunocompromised individuals following exposure to aquatic environments or healthy individuals that survive near drowning events [64–66]. Although human cases have been reported from multiple countries and US states [64–67], we know of no human cases reported from Texas. The association of human infections with water, particularly salt water or brackish water, as well as most environmental isolates being obtained from water [2, 64, 68–70], have led to the suggestion that *F. philomiragia* may be adapted for aquatic environments [64]. *F. philomiragia* was previously thought to have been isolated from sea water near Houston [7], but those isolates (TX07-7308, TX07-7310) have now been assigned to other *Francisella* spp. [2]. However, at least two human cases (one from the US state of Indiana and the other from Malaysia), notably both involving pneumonia, which is also common with other human *F. philomiragia* infections [65, 66], had no suspected water exposures [67, 71]. And a previous study [6] initiated following the detection of *Francisella* spp. DNA in aerosol samples collected in Houston in 2003 generated an *sdhA* sequence (0.36.C01) from a soil sample collected in Houston consistent with *F. philomiragia*. Although the taxonomy of *Francisella* has changed since that previous study was published [2], our analysis of that previous *sdhA* sequence, together with *sdhA* sequences from known isolates of *F. philomiragia* (Fig 3), confirms that the sequence from the previous Houston soil sample is consistent with *F. philomiragia*, strongly suggesting this species occurs in soil in the Houston area. This, together with our identification of *F. philomiragia* DNA on three separate air filters, suggests *F. philomiragia* occurs in and is aerosolized from soil in Houston. Of note, this species is able to infect mice and is virulent to them via intranasal delivery [72].

It was not possible to identify known *Francisellaceae* species from many of the 43 air filters that we examined (S3 Table) even though previous DNA extractions and PCR testing that was not part of this study identified the presence of *Francisella* spp. DNA on these filters and, following two rounds of enrichment, we generated sequencing reads from all 43 samples mapping to at least some of the 188,431 *Francisellaceae* probes (S4 Table). The presence of *sdhA* gene sequences was useful for identifying the likely presence of specific *Francisella* spp. present in some of these samples but we did not identify *sdhA* sequences from all 43 samples (S3 Table), likely because we did not include this genomic region in our probes as it also is found in other non-*Francisellaceae* bacterial species and we limited our probes to genomic regions exclusive to *Francisellaceae* species. Despite this, we identified *Francisella sdhA* sequences in 19 of the 43 air filter samples, likely due to by-catch. In addition, given the complex nature of environmental DNA captured on air filters [73], we elected to be conservative in our approaches for identifying MAGs for known *Francisellaceae* species and only included samples in species-specific phylogenies if they yielded MAGs. That said, another reason why we may not have identified known *Francisellaceae* species from some of the air filters is because they contain novel *Francisellaceae* species that were not included in the bait design. Although some genomic regions in novel species could share homology with a subset of our probes, the fragmented nature of these regions would likely prevent assembly of MAGs and subsequent analyses.

Sequence reads generated from four enriched samples collected in Houston (Air07, Air10, Air13, Air14) and all six enriched samples collected in Miami (Air38-43) mapped to <3% of the 188,431 *Francisellaceae* probes (range: 0.50-2.92%; S3 Table) and mapped to <4% of the probes specific to any one *Francisellaceae* species (range: 0.00-3.26%; S4 Table). However, sequence reads from all ten of these samples mapped to some of the probes specific to Clade 1 and/or Clade 2 (S4 Table) in the *Francisellaceae* global phylogeny. Indeed, two of the samples collected in Miami, Air40 and Air43, yielded reads that mapped to >37% of the 324 probes specific to clade 1 but did not yield *Francisella* MAGs. This may be because these two air filters – and perhaps others that did not yield *Francisella* MAGs – contained DNA from novel *Francisella* species that could not have been included in our bait capture system and, therefore, were not effectively captured and enriched by our approach. If these are novel species, the presence of reads mapping to Clade 1-specific probes in these samples suggest these unknown species may be closely related to *F. tularensis* as it also belongs to Clade 1 within *Francisella*.

Overall, our analyses of these air filters document that multiple known, and possibly some unknown, *Francisella* species can be, and possibly are regularly, aerosolized; whether these bacteria remained viable during and/or following aerosolization is currently unknown but *F. tularensis* has this capability [57]. In addition, these findings demonstrate the utility of utilizing an enrichment approach to increase the genomic signal of target species captured by air filters. This then facilitates detailed analyses of the target species, including assembly of MAGs in some cases, which can be difficult to accomplish with data generated via metagenomics sequencing of the original complex DNA extracts obtained from air filters.

#### Tick samples containing FLEs

Sequencing of the five complex DNA extracts from whole ticks/tick cell lines, following two rounds of capture and enrichment, produced an average of 6,925,142 (range: 413,912-12,086,278) sequence reads per sample, with an average of 80% of the reads classified as *Francisellaceae* (range: 67.6-93.9%; S3 Table). Reads from all five samples mapped to >72% of the 759 FLE-specific probes (S4 Table) and grouped into one *Francisella* MAG per sample, which ranged in size from 1.21-1.45Mb (S3 Table); these five MAGs clustered with other FLE genomes in the dendrogram (S2 Fig). Note that some initial enrichment results for tick samples D14IT15.2 and D14IT20 have been previously described [20] but they are included here for completeness and because additional analyses were conducted on these samples in this study.

The *Francisella* reads generated from the five enriched complex tick samples facilitated high-resolution phylogenetic analyses of FLEs from different as well as the same tick species. Although we generated and included genomic data for FLEs from three additional tick species (*D. albipictus*, *D. variabilis*, *H. rufipes*), some of the general phylogenetic patterns (Fig 7) remain similar to those previously described [26, 27, 74]: all 14 FLE genomes group together in a monophyletic clade within the *Francisellaceae* phylogeny and belong to the genus *Francisella* (S2 Fig), the FLE in *A. arboreus* (*F. persica*) is basal to all other FLEs, FLEs from hard ticks (Family Ixodidae: *A. maculatum*, *D. albipictus*, *D. variabilis*, *H. asiaticum*, *H. marginatum*, *H. rufipes*) are interspersed with FLEs from soft ticks (Family Argasidae: *A. arboreus*, *O. moubata*), FLEs from eastern hemisphere ticks (*A. arboreus*, *H. asiaticum*, *H. marginatum*, *H. rufipes*, *O. moubata*) and FLEs from western hemisphere ticks (*A. maculatum*, *D. albipictus*, *D. variabilis*) primarily group separately, and multiple FLE representatives from the same tick species group together (Fig 7). And, in general, our overall results in terms of phylogenetic relationships among FLEs present in different tick species match those from an existing gene-based genotyping system for FLEs [74, 75], with the caveat that *D. albipictus* and *D. variabilis* have not been previously analyzed with that system. However, our targeted genome enrichment approach provides significantly increased resolution among FLE samples because we can use much more of the shared core genome of these samples; the FLE phylogeny in Fig 7 is based upon 53,377 shared positions among the five FLE samples enriched in this study and the nine previously published FLE genomes. This provided resolution among even very closely related FLEs. For example, based upon sequencing of a >1,000 bp amplicon of the bacterial 16S rRNA gene, previous analyses [19] assigned an identical sequence type (T1) to the FLEs in the two *D. variabilis* samples also examined in this study (D.v.0160, D.v.0228), but 310 SNPs were identified between them when comparing the *Francisella* reads generated from the enrichment of these samples. Similarly, a previous analysis of >3,800 bp of sequence data generated from amplicons from multiple genes found no differences between seven representatives of the FLE in *H. rufipes* [74] but a total of 1,922 SNPs were identified between the *Francisella* MAGs generated from the two *H. rufipes* samples enriched and sequenced in this study (D14IT15.2, D14IT20), which were collected on the same day at the same location from two different birds [20]. This increased genetic resolution, and the ability to generate targeted FLE genomic data directly from whole, wild-caught ticks, offers the opportunity for future detailed studies of the natural population structure of these FLEs within single tick species. Of note, all of the previously published FLE genomes included in our analyses involved metagenomic sequencing of complex DNA extracted from whole ticks or specific tick organs [3, 26–30], with the exception of *F. persica*, to date the only FLE to be isolated; its genome was generated from genomic DNA [25].

**Fig 7.**
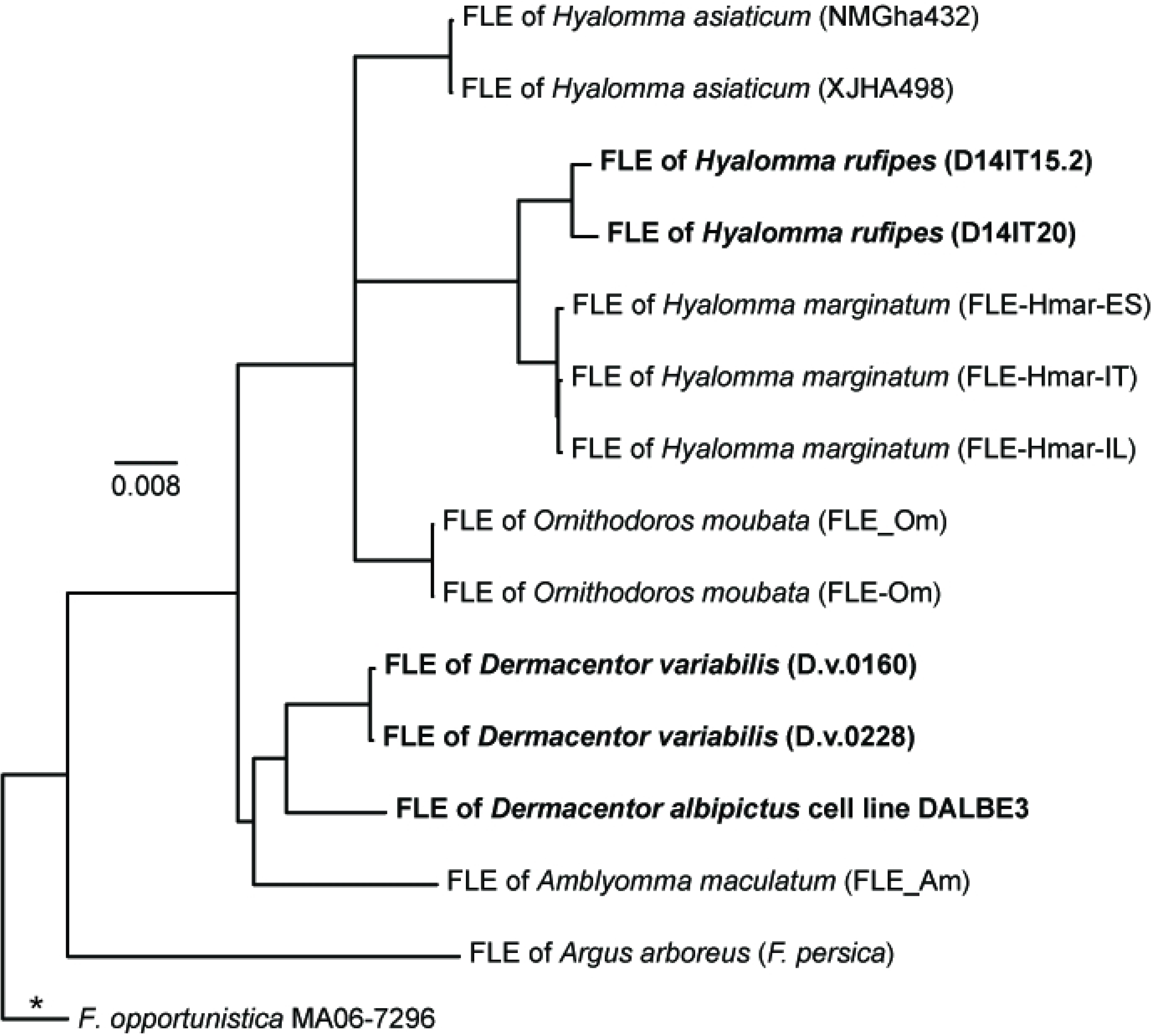
*Francisella*-like endosymbiont (FLE) phylogeny. Maximum-likelihood phylogeny based upon 53,377 core genome SNPs shared in nine publicly available FLE genomes (S1 Table) and DNA enriched from five whole ticks/tick cell lines (bold text). The phylogeny is rooted on the branch indicated with * and scale bar units are average nucleotide substitutions per site.

The *Francisella* MAGs assembled from the five enriched complex tick samples also allowed comparative analyses of gene content in FLEs. As previously noted for the published FLE genomes [3, 26, 27, 30], multiple CDSs in the FPI also are missing and/or disrupted in the genomes of the FLEs from *D. albipictus* (DALBE3), *D. variabilis* (D.v.0160, D.v.0228), and *H. rufipes* (D14IT15.2, D14IT20) generated in this study (S6 Table). However, the specific FPI genes that are missing or disrupted differs across all FLE genomes, even though intact versions of each of the 17 examined FPI CDSs are present in at least one of the 14 FLE genomes. This is consistent with the suggestions first put forward by Gerhart *et al* [3] that FLEs: 1) evolved from a pathogenic ancestor, 2) are relatively young, and, thus, 3) are in the initial stages of genome reduction. Loss and disruption of FPI CDSs in the FLE genomes is not surprising given that, in *F. tularensis* and *F. novicida*, these CDSs are associated with intracellular growth within vertebrate macrophages and virulence in mammalian hosts [62, 76, 77], conditions not encountered by FLEs. In contrast, all the examined biotin genes remain intact in the FLEs of *D. variabilis* (D.v.0160, D.v.0228) and *D. albipictus* (DALBE3), as well as in most of the other published FLE genomes (S7 Table), likely because this pathway allows FLEs to provide their ticks hosts with this essential B vitamin, as previously experimentally confirmed and/or suggested [26–28, 78]. Indeed, the only documented exceptions to this pattern to date are in the FLEs of *H. marginatum* (FLE-Hmar-ES, FLE-Hmar-IL, FLE-Hmar-IT) and *H. rufipes* (D14IT15.2, D14IT20), which are both involved in dual symbioses with *Midichloria* within their tick hosts; these *Midichloria* contain intact biotin pathways that are assumed to compensate for the disruption of this pathway in their associated FLE counterparts [20, 30]. Of note, previous attempts [20] to examine biotin gene content in metagenomic sequences of DNA extracts from samples D14IT15.2 and D14IT20 were unsuccessful due to low FLE sequence signal.

It was recently demonstrated that the *Coxiella*-like endosymbiont (CLE) of the of the Asian longhorned tick (*Haemaphysalis longicornis*), but not the tick itself, contains an intact shikimate pathway that synthesizes chorismate. *H. longicornis* then utilizes the chorismate produced by the CLE to produce serotonin, which influences its feeding behavior [79]. All the genes required for encoding the production of the enzymes in the shikimate pathway (*aroG*, *aroB*, *aroD*, *aroE*, *aroK*, *aroA*, *aroC*) are present and this pathway appears to be functional in *F. tularensis* [80, 81], and we found that homologs of these same genes are highly conserved in several FLE genomes (S8 Table). This suggests that some FLEs also may provide chorismate to their tick hosts for subsequent conversion to serotonin, but this would need to be experimentally confirmed. A notable exception to this pattern is the FLE in the *D. albipictus* DALBE3 cell line: multiple genes encoding the shikimate pathway are missing or disrupted in its genome. In addition, the MAG for the FLE of DALBE3 is significantly smaller than that of the other FLE genomes [30; S3 Table]; both patterns may reflect adaptation of this FLE to the more restricted niche of a tick cell line compared to a whole tick.

Symbiotic relationships between animal hosts and microorganisms are widespread and ancient [82]. These relationships are essential for obligate blood-feeding hosts, such as ticks, which require cofactors and vitamins not available in blood but that can be provided by their nutritional endosymbionts [78]. Key to understanding these relationships has been the generation of genomic data for both hosts and endosymbionts [82] but this can be obstructed by a lack of culturability of endosymbionts [83], which is the case for all FLEs except *F. persica*. Metagenomics analysis of whole or partial ticks containing FLEs has yielded important insights [3, 26–30], and metagenomics also has the added benefit of generating sequence data for other microorganisms within the tick and/or the tick itself. However, metagenomics analysis requires significant sequencing resources and, therefore, may be cost-prohibitive for some applications, especially if the emphasis of the study is only the FLE or some other endosymbiont. Our results demonstrate that the DNA capture and enrichment approach described here also can be used to generate robust genomics data for FLEs, even for FLEs not included in our probe design. Only FLEs from the tick species *A. arboreus*, *A. maculatum*, and *O. moubata* were utilized for the design of our enrichment system, yet this system still captured and enriched high-quality genomic DNA from FLEs present in other tick species, including *D. albipictus*, *D. variabilis*, and *H. rufipes*. As was previously noted for an enrichment system targeting *Wolbachia* endosymbionts [84], this finding demonstrates that the enrichment approach described here works across large evolutionary distances among FLEs.

### Recommendations for enrichment, sequencing, and analysis

The results of this study demonstrate the efficiency of DNA capture and enrichment from low signal samples containing *Francisellaceae* DNA. To maximize the signal for subsequent whole genome analyses, we recommend two rounds of enrichment when working with complex samples. In lower complexity samples and/or samples with a high starting proportion of *F. tularensis* or other target *Francisellaceae* DNA, a single enrichment may provide enough actionable information, depending on the nature of the investigation. We performed deep sequencing on enriched samples (average=9.1M reads, S3 Table) but this may be unnecessary for all applications. To determine the minimum number of reads required for accurate genotyping, we subsampled sequencing datasets generated from the eight enriched clinical samples from Turkey containing *F. tularensis* and placed the *F. tularensis* DNA present in those samples into a *F. tularensis* phylogeny using *WG-FAST*. The results (S3 Fig) demonstrate that an average of just 3000 paired-end Illumina reads (150nts long) could accurately place the *F. tularensis* present in those samples into the same phylogenetic positions as obtained with the full dataset (Fig 2). For some samples, just 1000 paired-end, 150nts reads could accurately genotype an enriched sample, providing strain level resolution needed for source attribution and contact tracing. This demonstrates that if a complex sample is efficiently enriched, then even shallow sequencing can be sufficient for strain level genotyping resolution.

Other applications could change the sequencing and enrichment strategy. For example, two enrichments and deep sequencing may be needed for more complete genome assemblies. For the complex air samples, we determined that if >50% of the signal is *Francisella* following enrichment, then a metagenome can be assembled, but individual MAGs cannot always be reliably assembled. If <50% of the signal is *Francisella* following enrichment, the fragmented assemblies can be taxonomically assigned, but typically cannot be assembled into contigs. Thus, for lower signal samples, >2 enrichments per sample may improve sequence yield, but that possibility was not investigated in this study.

### Considerations for design of enrichment systems

Because we designed our enrichment system using genome sequences from all known *Francisellaceae* species [2], the system described here works well for capturing, enriching, and sequencing genomic DNA of diverse *Francisellaceae* species present in different types of complex backgrounds. However, we did not include the entire *Francisellaceae* pan genome in our capture-enrichment system; we excluded CDSs that also are present in non-*Francisellaceae* spp., including genes that could cross amplify non-target bacteria (*e.g.*, rRNA genes). We excluded these regions to focus capture, enrichment, and sequencing reagents on genomic regions known to be specific to *Francisellaceae* spp., as we wanted to create a system that could be used with very complex samples, such as air filters. These complex samples contain DNA from many diverse organisms [73, 85] that may contain conserved regions of the non-exclusive CDSs also found in *Francisellaceae* spp., which could lead to non-target enrichment [86]. However, it is important to point out that this design consideration is context dependent. If, for example, the primary interest is to enrich *Francisellaceae* DNA present in less complex samples, such as those obtained from human tularemia samples, then the entire *Francisellaceae* pan genome could be included in the enrichment system. And if the focus of the investigation is on a single *Francisellaceae* spp. such as *F. tularensis*, then the enrichment system could be reduced to target just the pan genome of that individual species. Similarly, if novel *Francisellaceae* species are discovered and contain novel genomic components not found in any currently known *Francisellaceae* species, those novel components can be added to subsequent versions of the enrichment system.

## Conclusions

Using the pan-genome of all known *Francisellaceae* species, we designed a system to enrich low-level *Francisellaceae* DNA present in complex backgrounds. To validate the system, we enriched, sequenced, and analyzed DNA from spiked complex control samples, clinical tularemia samples, environmental samples (*i.e.,* tissue from a squirrel spleen and whole ticks), and air filter samples. The *F. tularensis*-positive clinical samples enriched very effectively, providing almost complete genomic coverage, which facilitated strain level genotyping. Obtaining most of the genome also allowed for the identification of novel SNPs, as well as a profile of gene content. The benefits of this enrichments system include: even very low target DNA can be amplified; the system is culture-independent, reducing exposure for research and/or clinical personnel and allowing genomic information to be obtained from samples that do not yield isolates; and the resulting comprehensive data not only provides robust means to confirm the presence of a target species in a sample, but also can provide data useful for source attribution, which is important from a genomic epidemiology perspective. In some situations, this system can outperform metagenomic sequencing, as demonstrated with the clinical tularemia samples, which requires very deep sequencing to provide signal, and even then, the complete genome may be difficult to obtain.

From a diagnostic/detection perspective, our identification of multiple, redundant, specific probes allows for robust and unambiguous confirmation of the presence or absence of *F. tularensis* in a sample (the same principles apply to the other species/clades for which specific probes were identified in this study). Upon enrichment and sequencing, all the samples utilized in this study that were confirmed by other approaches to contain *F. tularensis* DNA also yielded robust coverage of all 68 *F. tularensis*-specific probes (S4 Table). However, if the pathogen signal is low in enriched samples and/or shallow sequencing depth is obtained for an enriched sample, it is possible that not all the *F. tularensis*-specific probes may be identified in the resulting analyses even if *F. tularensis* is present in the sample. In addition, as the enrichment system was designed using the known diversity of *Francisellaceae* species but new species in this family continue to be discovered, future enrichment and sequencing of complex samples containing currently unknown *Francisellaceae* species could reveal that these new species contain some of the 68 probes that are currently thought to be specific to *F. tularensis*. Indeed, we identified partial coverage of several of the 68 *F. tularensis*-specific probes in multiple air filter samples (S4 Table). However, there was no coverage of the other *F. tularensis*-specific probes, which allowed us to confidently confirm that *F. tularensis* was not present in these samples. Thus, the redundancy provided by multiple specific probes means this system will remain a robust approach for confirmation of the presence or absence of *F. tularensis* in a sample even if a sample is low-quality and/or contains novel *Francisellaceae* species that contain a subset of these probes.

## Acknowledgements

We thank Timothy Kurtti and Ulrike Munderloh, University of Minnesota, for permission to use the DALBE3 tick cell line and Kailee Savage for assistance with laboratory work.

## Supporting information

**S1 Fig. Enriched dust sample.** Percentage of reads classified as *Francisellaceae* based on Kraken2 for the spiked and unenriched dust sample, and the same sample after one and two enrichments.

**S2 Fig. MAG dendrogram.** A cluster dendrogram of pairwise MASH distances. The cluster was generated from distances with Skbio.

**S3 Fig. *WG-FAST* phylogeny.** The reference phylogeny was inferred from all SNPs identified from a reference set of *F. tularensis* genomes (Table S1). The query samples were inserted into the phylogeny with *WG-FAST* using all called SNPs. The query samples are shown in red and the number at the end of each sample name indicates the number of randomly sampled paired end reads.

**S1 Table. List of genomes utilized in this study.** Accession and other information for all genomes utilized in this study.

**S2 Table. Master list of probes utilized in this study.** The name and sequence of all probes in the enrichment design. Annotation was provided for specific probes based on their specificity to individual species or clades.

**S3 Table. Complex samples enriched in this study.** Metadata for samples enriched and analyzed in this study.

**S4 Table. Probe coverage across enriched samples**. The breadth of coverage at 3x depth for samples across selected probes.

**S5 Table. FPI gene screening results for *F. tularensis***. Breadth of coverage at 3x depth for all enriched samples containing *F. tularensis* against locus tags in *F. tularensis*. Annotation was provided if a premature stop codon was gained, if there was a frameshift mutation, if a truncation in the gene was observed, or if the gene appeared to be intact, all based on Snippy annotations.

**S6 Table. FPI gene screening results for *Francisella*-like endosymbionts**. Breadth of coverage at 3x depth for all FLE samples against locus tags in *F. persica*. Annotation was provided if a premature stop codon was gained, if there was a frameshift mutation, if a truncation in the gene was observed, or if the gene appeared to be intact, all based on Snippy annotations. **S7 Table. Biotin gene screening results.** Breadth of coverage at 3x depth across genes associated with the biotin synthesis cluster in *F. persica*. Annotation was provided if a premature stop codon was gained, if there was a frameshift mutation, or if the gene appeared to be intact, all based on Snippy annotations.

**S8 Table. Shikimate gene screening results.** Breadth of coverage at 3x depth across genes associated with the shikimate synthesis cluster in *F. persica*. Annotation was provided if a frameshift mutation was observed or if the gene appeared to be intact, all based on Snippy annotations.

## Notes

### Competing Interest Statement

The authors have declared no competing interest.

